# Controlling confined collective organisation with taxis

**DOI:** 10.1101/2023.12.05.570159

**Authors:** Albane Théry, Alexander Chamolly, Eric Lauga

## Abstract

Biased locomotion is a common feature of microorganisms, but little is known about its impact on self-organisation. Inspired by recent experiments showing a transition to large-scale flows, we study theoretically the dynamics of magnetotactic bacteria confined to a drop. We reveal two symmetry-breaking mechanisms (one local chiral and one global achiral) leading to self-organisation into global vortices and a net torque exerted on the drop. The collective behaviour is ultimately controlled by the swimmers’ microscopic chirality and, strikingly, the system can exhibit oscillations and memory-like features.

---

As it is for their macroscopic counterparts, taxis – the ability to move in response to environmental cues – is crucial to the life of motile microorganisms [1]; they can respond to gradients in chemicals [2], light [3], viscosity [4] or gravity [5]. In turn, the directed motion of individuals drives new collective dynamics, such as the wellknown bioconvection of gyrotactic or phototactic algae [3, 6] or clustering instabilities [7]. A bias in the swimmer dynamics can take the form of passive control of cell orientation. For example, magnetotactic bacteria (MTB) exhibit magnetic moments and align to external fields [8]. In suspensions, this leads to pearling instabilities [9, 10], boundary-mediated clustering [11–13] and plume formation [14]. A consistent feature of biased collective dynamics is the prominence of confinement-mediated interactions; oriented swimmers tend to accumulate at boundaries and create dense regions where self-organisation occurs [14–16]. Given its significant potential for biological and biomedical control, a predictive framework for such self-organisation is needed.

Striking recent experiments showed that MTB can set in motion droplets 40 times larger than individual cells [17]. Specifically, suspensions of MTB inside waterin-oil droplets under a horizontal external magnetic field **B** = *B***ŷ** (Fig. 1a) spontaneously self-organise in a single large-scale vortex, thereby rotating the whole drop around the vertical *z*-axis. While global vortical flows are a staple of collective motion in circular or spherical confinement without director fields [18–20], in particular for bacteria [21–25], the vortex rotation direction is usually random [26]. Not so for MTB, where the vortex is consistently oriented in the positive *z*-direction [17], termed clockwise (CW), irrespective of the sign of *B*, hinting at a different mechanism for the onset of large-scale motion. Surprisingly, the vortex reverses to a negative *z*-rotation when the field is reversed to −*B***ŷ** [17]: a seemingly identical system now exhibits a different, counter-clockwise (CCW) bias. The collective behaviour therefore depends on the history of actuation, despite an underlying linear and inertialess flow.

**FIG. 1.**
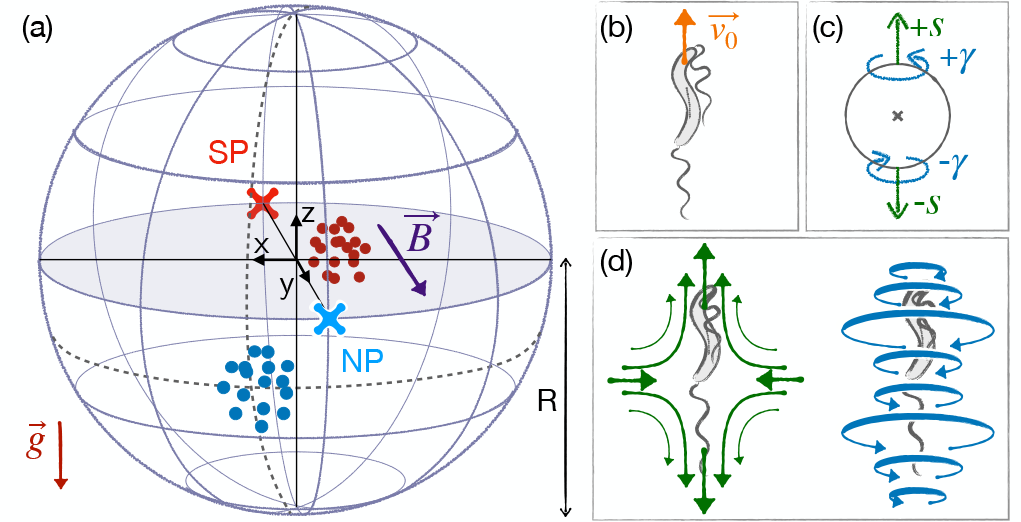
(a) Magnetotactic bacteria accumulate at the North (NP) and South Pole (SP) of the spherical drop. (b) Sketches of an individual MTB, (c) its hydrodynamic singularity model as dipoles of forces (*s*) and torques (*γ*), and (d) corresponding flow fields.

Here, we uncover the path to symmetry breaking and vortex formation in suspensions of biased swimmers under spherical confinement. Using simulations featuring long-range hydrodynamic interactions between cells with a chiral propulsive component, steric repulsion, and sedimentation, we identify two competing mechanisms for the onset of global rotation in the suspension. One, denoted (G), is global and stems from the interaction between two achiral populations at opposite poles of the drop, while the second is local, (L), and displayed by a single population of chiral swimmers gathering at one pole. Both mechanisms are explained through a minimal mathematical model involving hydrodynamic singularities. Notably, the local mechanism has a preferred rotation direction stemming from the swimmers’ chirality and leads to systematic CW rotation, as in Ref. [17]. This symmetry-breaking stems from the interaction of biased swimmers with an inclined (droplet) surface and is expected to be observed in other, non-spherical geometries. We also predict the onset of oscillatory flows for stronger, but physically relevant, chiralities. Moreover, the subtle competition between the two mechanisms controls the flow direction upon field reversal. Because of its sensitivity to chirality, this setup can be used to estimate and compare the chirality of biased microswimmers, a characteristic otherwise difficult to evaluate experimentally.

## Minimal model

We show that symmetry breaking is a physical rather than biological phenomenon with a minimal model for a suspension of *N*_*s*_ biased swimmers (Fig. 1), set up as follows. Each swimmer is described by its position and orientation. Initially, they are homogeneously distributed with random orientations within a sphere of radius *R* filled with a fluid of viscosity *μ*. Their dynamics are then set by (i) their alignment to the external magnetic field, (ii) self-propulsion, (iii) sedimentation, (iv) noise, and (v) steric and (vi) hydrodynamic interactions with other swimmers and the droplet boundary.

In detail, (i) a globally preferred orientation is set by *B***ŷ**, taking the role of an external director mechanism. The suspension is evenly split into north (NS) and south-seeking (SS) populations, which experience alignment torques towards the North (NP) and South pole (SP) respectively (Fig. 1a). The alignment strength is set by the field *B* and the magnetic moment normalised by the rotational drag coefficient *m*∼ . The bacteria also (ii) swim at a constant speed *v*_0_ and (iii) sediment due to a slight density difference with the medium 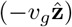. (iv) Thermal and active noise are included through Brownian noise in translation (diffusivity *D*_*t*_) and rotation (*D*_*r*_). (v) Hard-sphere steric interactions enforce a repulsion between nearby swimmers and keep them inside the sphere. (vi) Hydrodynamic interactions occur with the droplet surface, as well as with the flow from surrounding swimmers, themselves modified by the presence of a boundary.

In the dilute limit, hydrodynamic interactions are governed by the leading-order flow signature of the magnetotactic bacteria MSR-1 (Fig. 1d), which is an extensile force dipole (strength *s*) since cells are pushed by their aft helical flagella [27, 28]. Additionally, the flagellar rotation and associated body counter-rotation induce localised hydrodynamic torques, modelled as a rotlet dipole (strength *γ*), which decays faster than the stresslet but is the leading-order chiral signature. To explain the symmetry breaking, both flow singularities turn out to be necessary. To enforce the no-slip boundary condition on the droplet surface exactly, we use the method of images. We incorporate a perturbation to the bulk flow resulting from additional hydrodynamic singularities outside the sphere to match the flow due to the force [29, 30] and torque [30, 31] dipoles on the boundary [32]. Each individual swimmer is then advected and reoriented by the flow from its own image and the sum of bulk and image flows from other cells.

We scale lengths by the cell size *d*, time by *d/v*_0_, forces by *μv*_0_*d* and magnetic field strengths by *v*_0_*/*(*m*∼ *d*), and use dimensionless quantities; setup parameters are taken from Ref [17]. Values specific to the bacteria are drawn from experiments with MSR-1 when available [28, 33, 34] and we substitute unknown values of *s* and *γ* by those for *E. coli* [35, 36]. The supplementary material contains details of the numerical integration and the influence of all parameters.

### Results

Our simulations reveal that the magnetic field strength, *B*, acts as the control parameter for a transition from disorder without director field to a self-organised state where the ± *x*-symmetry is broken (Fig. 2a). For weak fields, the NS and SS cell populations separate and accumulate at their preferred pole, but the system remains symmetric in ± *x* (Fig. 2a(ii)). As |*B*| increases, the clusters move azimuthally to an asymmetric steady-state configuration, displaced clockwise from the magnetic poles (Fig. 2a(iii)-(iv) and b). As a result of this shift in position, a global CW vortex emerges in the *x*-*y* plane, accompanied by two narrow recirculation regions between each cluster and the boundary, in agreement with experimental observations [17].

**FIG. 2.**
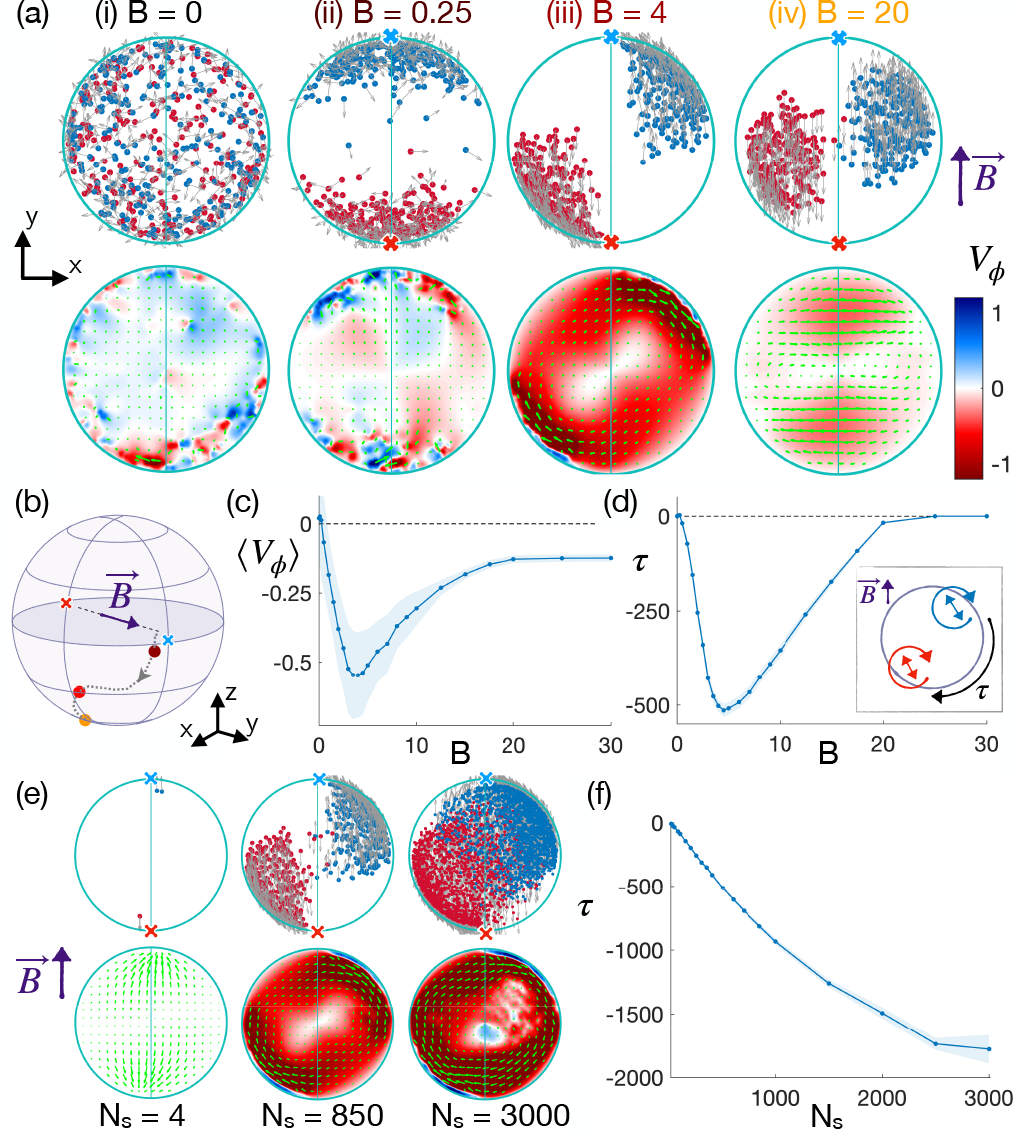
symmetry breaking and flow transition under an external field. (a) Snapshots for different magnetic field strengths. Top: position of the NS (blue) and SS (red) swimmers. Bottom: flow in the equatorial plane, in-plane velocity vectors (green), superposed with a density plot of the azimuthal component, *V*_*ϕ*_. (i) Disordered system at *B* = 0. (ii) Clusters form at the poles when the field is turned on. (iii, iv) Clusters slide CW at higher fields, creating a global CW vortex in the *x*-*y* plane. (b) Average position of the above NS clusters. (c) Mean value of spatially averaged vortex strength, ⟨*V*_*ϕ*_⟩, with shaded steady-state standard deviation. The CW vortex is strongest at *B* = *B*^*∗*^ ∼ 4. (d) Total magnetic torque, *τ*, as a function of *B*. Inset: sketch of the torque generation mechanism, which vanishes when *B* = 0 or *B >* 20. (e) Phenomenology for different concentrations: (*N*= 4) Broken symmetry for just two swimmers from each population, (*N*= 850) emergence of a vortex, and (*N*= 3000) partial mixing at high concentrations (f) Torque on the drop as a function of *N*.

The average azimuthal component of the fluid velocity in the equatorial plane, ⟨*V*_*ϕ*_⟩ (Fig. 2c), quantifies the transition to a vortical flow. It exhibits a maximum at a strong but intermediate value of the dimensionless field *B*^*∗*^ ∼ 4 (above the range of Ref. [17]). This optimum stems from a competition between alignment with the boundary and with the field. Partial alignment with the boundary at weak fields (Fig. 2a(iii)) promotes upward swimming against gravity and a greater separation between clusters, leading to stronger circulation. At higher field strengths, symmetry is still broken but the vortex generated by the sedimented swimmers is weaker (Fig. 2a(iv)). The misalignment of swimmers with the field at the boundary also results in a net magnetic torque which is predominantly CW in both NS and SS cell clusters (Fig. 2d, inset). This net torque (*τ* ) is transmitted to the droplet, which rotates if freely suspended, as seen experimentally [17], with an optimum close to *B*^*∗*^ (Fig. 2d).

Notably, symmetry breaking occurs for as few as two swimmers from each population (Fig. 2e). The vortex and torque then increase approximately linearly with the number of swimmers *N*_*s*_. For denser suspensions, the vortex destabilises the clusters into a mixed spinning core, and the torque *τ* plateaus at *τ*_*p*_ ≈ 0.6 nN μm, consistently with experimental values *τ* ≈ 0.2 − 0.5 nN μm (Fig. 2f).

Our simulations reproduce the experiments of Ref. [17], showing that hydrodynamic interactions are at the essence of the observed symmetry breaking. Beyond this, our simulations allow us to explore a range of microscopic parameters to explain its underlying mechanisms. Notably, we study achiral bacteria (i.e. no torque dipole, *γ* = 0), and find that rotational symmetry is only broken if both NS and SS populations coexist. Even then, the system randomly develops either a CW or a CCW vortex, unlike the consistent CW pattern in experiments.

In contrast, for moderately chiral bacteria (*γ >* 0), symmetry breaking occurs CW systematically for either one (for ex, just NS) and two (NS and SS) populations. Remarkably, when either chirality is strong (large *γ*), or the force dipole *s* and gravity *v*_*g*_ are weak, the stationary state vanishes and gives way to oscillations. The clusters precess around the poles, and the flow alternates between CW and CCW.

This suggests the presence of two distinct mechanisms causing the observed vortex: a global achiral one (G) in which NS and SS clusters interact at long range, and a local chiral one (L) that allows a single population to break symmetry.

### (G) Global symmetry breaking

The first mechanism, achiral interactions between NS and SS clusters, involves only the leading-order force dipole signature of the swimmers. As shown in Figs. 1d and 3a, extensile force dipoles (*s*) create a lateral attractive flow [37]. For aligned pusher swimmers parallel to one another, in particular for bacteria swimming perpendicular to a plane wall, this lateral flows results in a well-known attraction [38] and clustering [11, 14]. As a result, for sufficiently strong fields the swimmers from a population in the drop align, attract, and form clusters that move cohesively on the boundary. For two populations, each cluster pushes flow away from its pole and thus destabilises the centre of mass of the cluster at the opposite pole. This configuration is unstable. Both clusters then move laterally, leading to broken symmetry and a vortical flow. Gravity acts as a stabiliser in the *z*-direction, so the instability occurs horizontally. Crucially, for achiral swimmers symmetry is broken randomly, and both CW and CCW vortices emerge with equal probability (Fig. 3b).

**FIG. 3.**
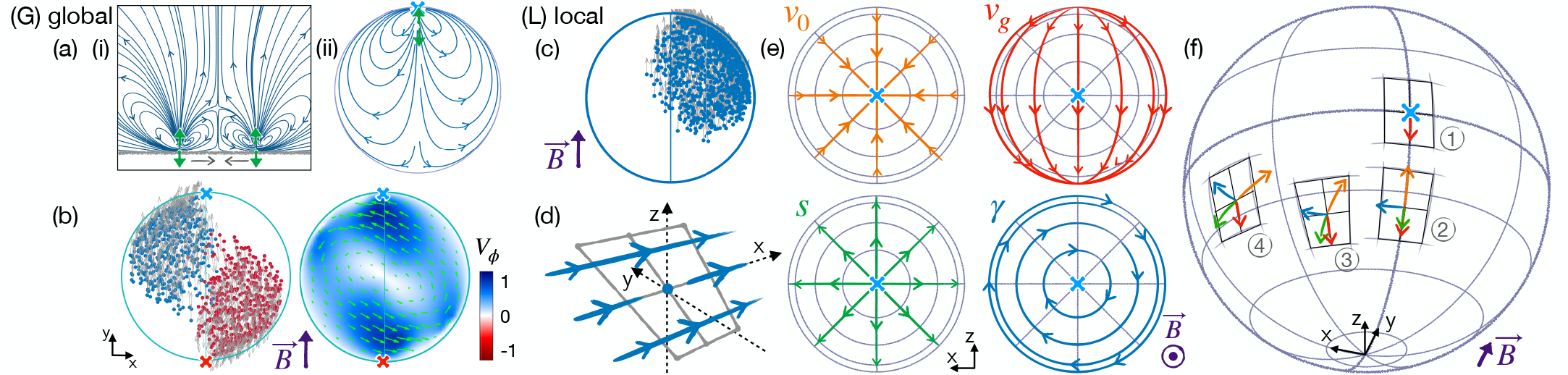
Global (G) and Local (L) mechanisms for symmetry breaking. (a) (i) Streamlines from two force dipoles perpendicular to a wall. The inward lateral flow results in an effective attraction (grey arrows). (ii) Streamlines from a force dipole at the NP. For a cluster at the SP, this flow is destabilising. (b) (G)-symmetry breaking for two achiral (*γ* = 0 populations, with no preferred CW/CCW direction. (c) Directed (L) CW symmetry breaking for a single chiral population. (d) Net flow from a dipole of torques *γ* along **ŷ** near a no-slip inclined boundary. (e) Contributions to the NS cluster displacement: self-propulsion *v*0, gravity *v*, force dipoles *s* (attraction to parallel walls), torque dipoles *γ* (CW precession around the NP). (f) Dynamic sequence of cluster positions during a (L) symmetry breaking, from the NP (1) to the steady-state (4).

### (L) Local symmetry breaking

The systematic selection of CW rotation relies on a second physical mechanism. Our simulations reveal that it involves the higherorder effect of chirality *γ* and occurs even for a single population (Fig. 3c), hinting at the significance of local hydrodynamic interactions with the boundary. Under the assumption that the swimmers are aligned with the external field, a single population of, say NS, swimmers form a cluster, which moves cohesively along the droplet boundary. We analyse its trajectory by considering each contribution to the motion individually (Fig. 3e). (i) Self-propulsion along the +*y*-direction drives swimmers towards the magnetic NP. (ii) Gravity uniformly displaces the cluster downwards to 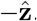. (iii) Extensile force dipoles are hydrodynamically attracted to parallel surfaces through their image. In our geometry, this drives them to the *y* = 0 plane, opposite to self-propulsion. These three contributions are ± *x*-symmetric and do not yet explain vortex formation. However, (iv) a torque dipole facing an inclined boundary creates a net lateral flow in its vicinity, oriented tangential to the boundary and orthogonal to the swimmer axis (Fig. 3d). While no individual swimmer can advect itself in this way, the collective action of bacteria on each other laterally displaces the cluster as a whole, while cohesion is maintained by swimmer-swimmer attraction. On the drop surface, this lateral translation results in a precession around the *y*axis.

A steady-state position for the cluster then exists if these four effects balance locally, which can only occur at a shifted CW position (Fig. 3f). If *γ* (i.e. the contribution in (iv)) is too strong, there is no stationary point and the clusters precess around the poles, creating the oscillating regime we observe numerically.

Our simulations reveal therefore that the systematic CW vortex in experiments results from the combination of two distinct physical mechanisms: (G) a global achiral interaction between cell clusters leads to global broken symmetry with no preferred direction, and (L) a local chiral confinement-mediated mechanism leads to local CW symmetry breaking.

### Field reversal

The coexistence of two instabilities explains the puzzling experimental observation that the vortex direction is reversed upon reversal of the magnetic field, although it initially rotates CW regardless of its orientation [17]. In a uniform population, both instabilities cooperate in shifting the location of the clusters laterally, with the CW direction set by the local one. In contrast, when the field is reversed, it acts on swimmers that are already asymmetrically distributed. The chiral

CW instability (L) then competes with the populationpopulation repulsion (G) that keeps clusters on the side they were already on, and the new steady state depends on the dominant mechanism. Below a threshold chirality *γ* (which depends *B* and *N*_*s*_) the achiral populationpopulation hydrodynamic repulsion (G) keeps swimmers on the same (left or right) side of the *y* − *z* plane as they exchange poles: the flow is thus reversed to CCW, as in experiments (see Fig. 4(i-ii)). Above this threshold, the chiral hydrodynamic effect (L) is sufficient for the populations to shift CW azimuthally and exchange sides, so the flow and torque eventually return to CW (Fig. 4(iii)).

**FIG. 4.**
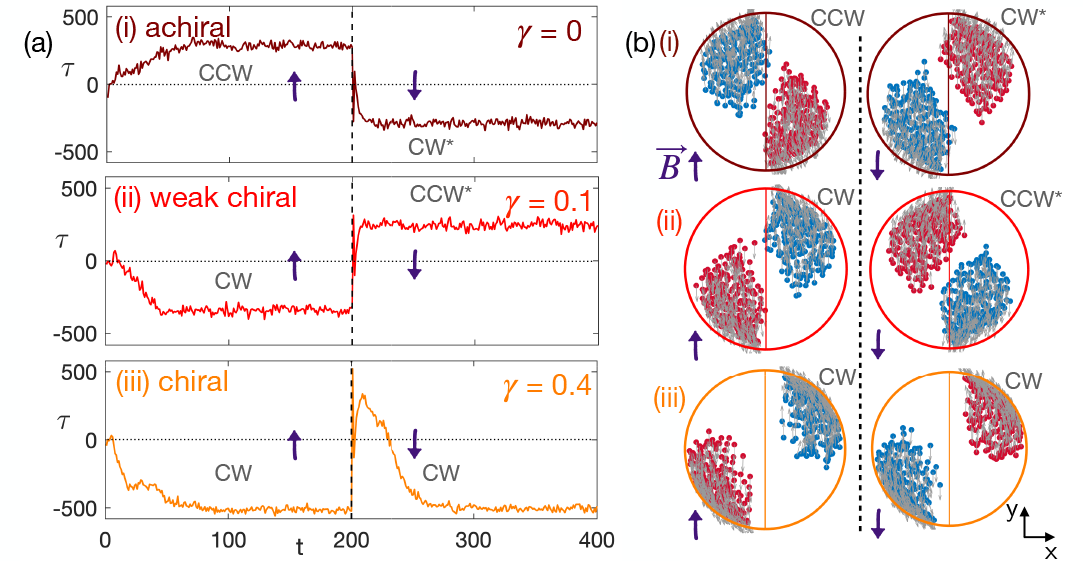
Magnetic field reversal in (i) achiral, (ii) weakly chiral, (iii) chiral suspensions. (a) The reversal at *t* = 200 (dashed line) causes the reversal of the torque *τ* (denoted as ^*∗*^) for achiral and weakly chiral swimmers (i, ii) while for a stronger chirality, the torque regains its initial CW direction (iii). (b) Stationary-state position of swimmers before (left) and after (right) the field reversal. (i) The achiral system is symmetric with respect to the (*x* − *z*) plane. (ii) The weakly chiral has a preferred CW direction, which is lost after the reversal because of the (G) interaction between the populations. (iii) Stronger chirality (L) overcomes this repulsion and the populations shift back to CW.

Reversing the field differentiates between swimmers where force propulsion dominates, such as MSR-1, and more chiral ones. Since increasing chirality further induces oscillations, this could be used as a proxy to estimate the relative importance of forces and torques in propulsion.This is particularly relevant because the degree of cellular chirality is difficult to access experimentally. To our knowledge, the dipole strengths for MSR-1 have not been measured.

Our fundamental model reveals how a sequence of symmetry-breaking controls the collective dynamics of confined biased swimmers and the emergence of a global vortex in the equatorial plane. Two distinct mechanisms can trigger collective motion, (G) globally through the long-range repulsion of NS and SS populations and (L) locally for chiral swimmers. In the latter, a few swimmers of a single population are enough to break the lateral symmetry because of hydrodynamic interactions with inclined walls. This tunable and externally controlled collective motion offers a promising framework for the analysis of populations of biased cells. The reversal dynamics and oscillations capture the underlying microscopic properties of the swimmers, in particular the relative strength of hydrodynamic forces and torques, which may otherwise be hard to measure. For physically relevant values, oscillations arise in the steady state of strongly chiral swimmers, while the vortex reversal upon field reversal seen experimentally in [17] is identified as the signature of weak chirality. The rotation direction then depends on the history of the suspension even in a linear system.

This project has received funding from the European Research Council under the European Union’s Horizon 2020 research and innovation program (Grant No. 682754 to E.L.). A.T. received support from the Simons Foundation through the Math + X grant awarded to the University of Pennsylvania.

