## Supplementary information for "Controlling confined collective organisation with taxis"

### Supplementary Material
### Controlling confined collective organization with taxis.

Albane Théry,[1, 2, *] Alexander Chamolly,[3, 4, †] and Eric Lauga[1, ‡]

[1] *Department of Applied Mathematics and Theoretical Physics, University of Cambridge, Cambridge CB3 0WA, UK*
[2] *Department of Mathematics, University of Pennsylvania, 19104 Philadelphia PA, US*
[3] *Institut Pasteur, Université de Paris, CNRS UMR3738, Developmental and Stem Cell Biology Department, F-75015 Paris, France*
[4] *Laboratoire de Physique de l'Ecole normale supérieure, ENS, Université PSL, CNRS, Sorbonne Université, Université de Paris, F-75005 Paris, France*

(Dated: December 5, 2023)

#### I. DETAILS OF THE SIMULATIONS

##### A. Parameters

The value of each parameter used in the simulations when not stated otherwise is given in Table I. Upon non-dimensionalisation, the unit length, time, strength and magnetic field are $4\mu$m, 0.2s, 0.08pN and 1.14mT respectively.

For computational feasibility, the number of bacteria is taken here to be at most $N_s = 3000$.

TABLE I. Parameters for the simulations, with experimental values [1] and their non-dimensional equivalent, as well as the default values used in simulations.

| Parameter | Experimental value | Non-dim. | Default |
|---|---|---|---|
| $N_s$ | $10^{17}$cells m$^{-3}$ | $10^4$ | 500 |
| $n$ | $N_s/2$ | $N_s/2$ | 250 |
| $B$ | $0.1 - 4$mT | 0.11 - 4.5 | 5 |
| $R$ | $20 - 120$¯m | 5 - 30 | 10 |
| $v_0$ | $20 - 40$¯m s$^{-1}$ | 1 | 1 |
| $v_g$ | 3¯m s$^{-1}$ | 0.15 | 0.15 |
| $\tilde{m}\mu$ | 57nN ¯m$^{-2}$ mT$^{-1}$ [2, 3] | | |
| $s$ | 0.57pN [4] | 7 | 7 |
| $\gamma$ | 0.35pN ¯m [4] | 1.1 | 0.4 |
| $\mu$ | $10^{-3}$pN s ¯m$^{-2}$ | 1 | 1 |
| $d$ | 4¯m [3, 4] | 1 | 1 |
| $D_r$ | 0.25rad$^2$ s$^{-1}$[5] | 0.05 | 0 |
| $D_t$ | 200¯m$^2$ s$^{-1}$ [6] | 2.5 | 0.1 |

##### B. Components

Initially the $N_s$ bacteria are positioned uniformly in the sphere at $\boldsymbol{R} = (\boldsymbol{r}_i)$, with random unit orientations $\boldsymbol{P} = (\boldsymbol{p}_i)$ [7].

The evolution of position and orientation is of swimmer $(i)$ is given by

$$\dot{\boldsymbol{r}}_i = v_0\boldsymbol{p}_i - v_g\hat{\boldsymbol{z}} + \boldsymbol{v}_i^{\text{im}} + \boldsymbol{v}_{\text{wall}\to i}^{\text{ste}} + \sum_{j\neq i}\left(\boldsymbol{v}_{j\to i}^{\text{hyd}} + \boldsymbol{v}_{j\to i}^{\text{ste}}\right) + \sqrt{2D_t}\,\boldsymbol{\xi}_t, \tag{1}$$

$$\dot{\boldsymbol{p}}_i = \left[\boldsymbol{\omega}_i^B + \boldsymbol{\omega}_i^{\text{im}} + \sum_{j\neq i}\boldsymbol{\omega}_{j\to i}^{\text{hyd}} + \sqrt{2D_r}\,\boldsymbol{\xi}_r\right]\times\boldsymbol{p}_i. \tag{2}$$

With the contributions to the speed:

*
†
‡

- constant swimming speed: $v_0 \boldsymbol{p}_i$,
- sedimentation speed $v_g \hat{\boldsymbol{z}}$,
- hydrodynamic speed $\boldsymbol{v}_i^{\text{hyd}}$ from other bacteria $\boldsymbol{v}_{j\to i}$ and all the images $\boldsymbol{v}^*_{j\to i}$,
- steric hard core interactions with the bacteria $\boldsymbol{v}^{\text{ste}}_{j\to i}$ with strength controlled by $d_{\text{ste}} = d$,
- steric interactions with the wall $\boldsymbol{v}^{\text{ste}}_{\text{wall}\to i}$ to ensure that $|\boldsymbol{r}_i|< R - d/2$
- noise with diffusion coefficient $D_t$ in translation, with with $\boldsymbol{\xi}_t$ a Gaussian white noise,

and to the rotation speed:

- hydrodynamic reorientation from other bacteria $\boldsymbol{\omega}^{\text{hyd}}_{j\to i}$ and all the images $\boldsymbol{\omega}^{(h*)}_{j\to i}$,
- noise with diffusion coefficient $D_r$ in rotation, with with $\boldsymbol{\xi}_r$ a Gaussian white noise,
- magnetic reorientation.

At each time step d$t$, we compute the $(3 \times N_s)$ matrices for velocities $\boldsymbol{U} = (\boldsymbol{v}_i)$ and rotation velocity $\boldsymbol{\Omega} = (\boldsymbol{\omega}_i)$. The corresponding matrices for position and orientation, $\boldsymbol{R}$ and $\boldsymbol{P}$ are then updated with a first order Euler method

$$\begin{aligned}
&\boldsymbol{R}(t+\mathrm{d}t) = \boldsymbol{R}(t) + \mathrm{d}t\,\boldsymbol{U} + \sqrt{D_t \mathrm{d}t}\,\boldsymbol{\xi},\\
&\boldsymbol{P}(t+\mathrm{d}t) = \boldsymbol{P}(t) + \mathrm{d}t\,\boldsymbol{\Omega}\times\boldsymbol{P}(t) + \sqrt{D_r \mathrm{d}t}\,\boldsymbol{\xi}\times\boldsymbol{P}(t) \quad \text{and} \quad \boldsymbol{P}(t+\mathrm{d}t) = \boldsymbol{P}(t+\mathrm{d}t)/|\boldsymbol{P}(t+\mathrm{d}t)|.
\end{aligned} \tag{3}$$

**Hydrodynamic interactions.** Each swimmer is a pair of Stokeslets $s$ and of rotlets $\gamma$ at $\pm d/2$ from its centre $\boldsymbol{x}_i$. We also assume that the swimmers are spherical and much smaller than the velocity gradient length scale, so the advection velocity due to the flow of the swimmers in the no-slip sphere is equal to the flow speed at $\boldsymbol{x}_i$, namely

$$\boldsymbol{v}_i^{\text{hyd}} = \sum_{j\neq i} \boldsymbol{v}_{j\to i} + \sum_j \boldsymbol{v}^*_{j\to i}. \tag{4}$$

The flow from swimmer $j \neq i$ is the sum of the flows due to the two point-forces and two point-torques, which reads

$$\begin{aligned}
\boldsymbol{v}_{j\to i} =&\boldsymbol{v}^{(s)}\left(\boldsymbol{x}_i - (\boldsymbol{x}_j + d/2\boldsymbol{p}_j), s\boldsymbol{p}_j\right) + \boldsymbol{v}^{(s)}\left(\boldsymbol{x}_i - (\boldsymbol{x}_j - d/2\boldsymbol{p}_j), -s\boldsymbol{p}_j\right)\\
&+ \boldsymbol{v}^{(\gamma)}\left(\boldsymbol{x}_i - (\boldsymbol{x}_j + d/2\boldsymbol{p}_j), \gamma\boldsymbol{p}_j\right) + \boldsymbol{v}^{(\gamma)}\left(\boldsymbol{x}_i - (\boldsymbol{x}_j - d/2\boldsymbol{p}_j), -\gamma\boldsymbol{p}_j\right),
\end{aligned} \tag{5}$$

The velocity from a single stokeslet and rotlet are

$$\boldsymbol{v}^{(s)}(\boldsymbol{x},\boldsymbol{s}) = \frac{1}{8\pi\mu}\left[\frac{\boldsymbol{s}}{r} + \frac{(\boldsymbol{s}\cdot\boldsymbol{x})\boldsymbol{r}}{x^3}\right] \quad \text{and} \quad \boldsymbol{v}^{(\gamma)}(\boldsymbol{x},\boldsymbol{s}) = \frac{1}{8\pi\mu}\frac{\boldsymbol{\gamma}\times\boldsymbol{x}}{r^3} \tag{6}$$

respectively. The contribution from the hydrodynamic image is the sum of the individual images for the two stokeslets and the two rotlets. For the derivation, all the singularities are considered as the sum of an axisymmetric and a transverse component. The resulting flow for each term is found using results by Maul and Kim [8] for the axisymmetric Stokeslet, Shail [9] for the transverse Stokeslet, Chamolly and Lauga [10] for the axisymmetric rotlet and by Hackborn et al. [11] for the transverse rotlet. Their expressions are detailed in section III.

The reorientation speed is obtained similarly, assuming that the swimmers are spherical and much smaller than the length scale of the flow gradients

$$\boldsymbol{\omega}_i = \frac{1}{2}\left[\sum_{j\neq i}\boldsymbol{\omega}_{j\to i} + \sum_j \boldsymbol{\omega}^*_{j\to i}\right], \tag{7}$$

with the contribution from a given swimmer again stemming from the sum of the four hydrodynamic singularities

$$\begin{aligned}
\boldsymbol{\omega}_{j\to i} =&\boldsymbol{\omega}^{(s)}\left(\boldsymbol{x}_i - (\boldsymbol{x}_j + d/2\boldsymbol{p}_j), s\boldsymbol{p}_j\right) + \boldsymbol{\omega}^{(s)}\left(\boldsymbol{x}_i - (\boldsymbol{x}_j - d/2\boldsymbol{e}_j), -s\boldsymbol{p}_j\right)\\
&+ \boldsymbol{\omega}^{(\gamma)}\left(\boldsymbol{x}_i - (\boldsymbol{x}_j + d/2\boldsymbol{e}_j), r\boldsymbol{p}_j\right) + \boldsymbol{\omega}^{(\gamma)}\left(\boldsymbol{x}_i - (\boldsymbol{x}_j - d/2\boldsymbol{p}_j), -r\boldsymbol{p}_j\right).
\end{aligned} \tag{8}$$

The vorticities from the singularities are

$$\boldsymbol{\omega}^{(s)}(\boldsymbol{x},\boldsymbol{s}) = \frac{2}{8\pi\mu}\frac{\boldsymbol{s}\times\boldsymbol{x}}{r^3} \quad \text{and} \quad \boldsymbol{\omega}^{(\gamma)}(\boldsymbol{x},\boldsymbol{s}) = \frac{1}{8\pi\mu}\left[-\frac{\boldsymbol{\gamma}}{r^3} + \frac{3(\boldsymbol{\gamma}\cdot\boldsymbol{x})\boldsymbol{x}}{r^5}\right]. \tag{9}$$

As we did for the velocities above, we additionally derive exactly and include the image vorticities for each hydrodynamic singularity, with the expressions given in section III.

The solutions that we use for the hydrodynamic velocities and vorticities are singular at $\boldsymbol{x}_j$. To ensure that the local values of the flow velocities induced by swimming remain physically relevant close to a particle, we, therefore, set maximum values for the local velocity and vorticity generated by a swimmer. Measurement of the flow near a swimming *E. coli* by Drescher et al. [12] and detailed simulations [4] support the idea of a threshold velocity for the flow in the near field. We chose the threshold to be the maximum of the norm of the velocities and vorticities due to the pair of Stokeslets and rotlets, respectively, at a distance of $d_{\min}$ from the centre $\boldsymbol{x}_j$ of the swimmer. In our simulations, we take $d_{\min} = 1$. In the absence of a tractable regularisation function for stokeslets and rotlets inside a sphere, $d_{\min}$ acts as a proxy for regularisation as it prevents the velocities and vorticities from taking singular values.

The form of the hydrodynamic interactions used here is relevant for dilute suspensions. As confined magnetotactic bacteria tend to accumulate against boundaries, we expect this assumption not to hold in some regions of the sphere for high numbers of swimmers, comparable to the experimental concentration. In such high-density regions, we would expect the flow from the swimmers to be screened and hence weaker than in our simulations. We do not expect this restriction to affect the qualitative behaviour of the system, but the quantitative values of the simulated flows are likely to exceed experimental values. A possibility to get quantitative comparisons with experimental data could then be to rescale them to account for this screening.

**Steric interactions.** To avoid overlaps between the swimmers, we include a steric interaction between swimmers that are less than a radius away from each other, of the form

$$\boldsymbol{v}^{(ste)} = -\sum_{j\neq i} \delta_{|\boldsymbol{x}_i - \boldsymbol{x}_j|<d} \left(\frac{d_{\text{ste}}}{|\boldsymbol{x}_i - \boldsymbol{x}_j|}\right)^6 \frac{1}{|\boldsymbol{x}_i - \boldsymbol{x}_j|}(\boldsymbol{x}_i - \boldsymbol{x}_j). \tag{10}$$

The steric repulsion decays as $1/r^6$ and its strength is set by the typical lengthscale $d_{\text{ste}}$, with a default value of 0.2.

In addition, after each time step, we ensure that the swimmers stay inside the drop and at a minimal distance $d/2$ of the boundaries.

**Sedimentation.** The constant downward sedimentation speed $\boldsymbol{v}_g$ is obtained by considering the sedimentation of a sphere of diameter $d$ in the bulk of a viscous fluid. This gives $v_g = 2(d/2)^2 g\Delta\rho/(9\mu) \approx 3\,\text{\textasciimacron m}\,\text{s}^{-1}$ [13].

**Magnetic reorientation.** The magnetic torque reads

$$\boldsymbol{\omega}_m = \epsilon B \frac{m}{\zeta_s} \boldsymbol{p}_i \times \hat{\boldsymbol{y}}, \tag{11}$$

where $m$ is the magnetic moment and $\zeta_s$ is the drag on the swimmer for reorientation orthogonal to its long axis. Both were measured experimentally [2, 3], and we define $\tilde{m} = m/\zeta_s$.

We ensure that the magnetic field causes a reorientation at most aligned to the field axis $\hat{\boldsymbol{y}}$. If overshooting occurs and $|B\tilde{m}\boldsymbol{p}_i \times \hat{\boldsymbol{y}}| > \arccos(\boldsymbol{p}_i \cdot \hat{\boldsymbol{y}})$, we set instead that $\boldsymbol{p}_i(t + \mathrm{d}t) = \epsilon\hat{\boldsymbol{y}}$, so that the swimmer is aligned with the field and oriented towards its preferred pole.

**Noise.** We also include noise, both in translation and in orientation, with diffusion coefficient $D_t$ and $D_r$ chosen to be independent, as shown in Eq. (3) [14].

##### C. Implementation

There are also two parameters in our model that are not measured experimentally, the distance which sets the maximal hydrodynamic speed $d_{\min}/d$ and that for steric interactions $d_{\text{ste}}/d$.

In our simulations, we can tune macroscopic parameters, the number of swimmers, the magnetic field and the drop radius, and microscopic ones, the strength of gravity, the strength of the forces and torques due to the propulsion, the strength of steric interactions and the noise. We use a dimensionless time step $\mathrm{d}t = 0.05$ and an explicit Euler scheme to integrate the equations of motion.

#### II. INFLUENCE OF THE MICROSCOPIC PARAMETERS

##### A. Collective effects with one or two populations and achiral swimmers

We run simulations for different numbers of swimmers when taking into account one (NS) or two (both NS and SS) populations of swimmers in Fig. S1.

For a single NS population of achiral ($\gamma = 0$) swimmers, no vortex forms. If both NS and SS swimmers are present on the other hand, a vortex forms above a threshold number of swimmers. This vortex has no preferred CW-CCW direction. It also generates a torque on the droplet, which is plotted in Fig. S1 for varying concentrations of swimmers.

A vortex forms for a single NS population of chiral swimmers, but the resulting torque grows slower with $N_s$ than for two populations.

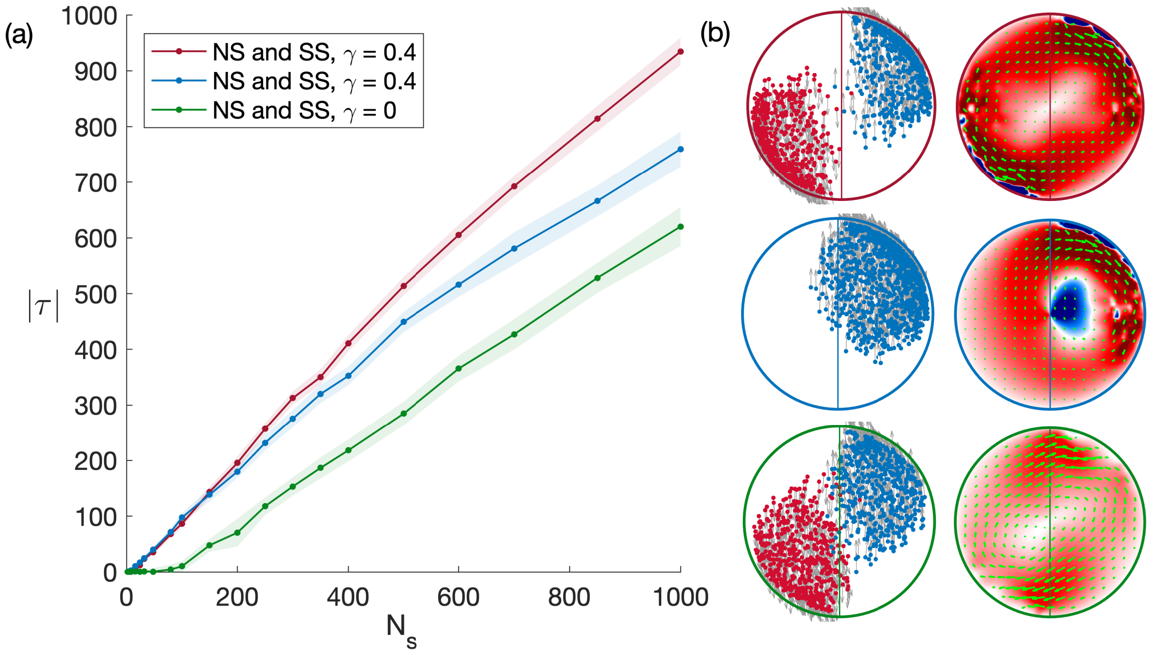


FIG. S1. Symmetry-breaking for two (red, top) or one (blue, centre) populations of chiral swimmers, and two populations of achiral swimmers (green, bottom) at $B = 5$. (a) Absolute value of the torque on the drop for $N_s$ swimmers, chiral or not, split into two populations or not, with the shaded region showing the standard deviation in time. (b) Position of the swimmers and equatorial flow for $N_s = 1000$. With a SS and a NS population, a symmetry breaking occurs and a torque arises for both achiral and chiral swimmers, but the chiral mechanism is both stronger and more robust at high concentrations. It also has a CW direction, while the achiral torque can be either CW or CCW. A torque is also created for a single chiral population but saturates for higher numbers of swimmers.

###### 1. *Interplay of force-based propulsion, gravity and chirality*

We vary the Stokeslet strength $s$, rotlet strength $\gamma$ and gravity $v_g$ while keeping constant the other two parameters, and plot the corresponding torque on the droplet and vortex strength in Fig. S2a. In all three cases, there is an optimal value for the collective motion which maximises both the average azimuthal velocity and the torque. When the system is less chiral than this, so for lower $\gamma$ or conversely higher $v_g$ or $s$, both vortex strength and torque decrease steadily. On the other hand, for very chiral swimmers, the stationary state of the system disappears. At long times, both clusters then undergo periodic trajectories around their pole, not always in phase, and the vortex flow alternates between CW and CCW directions. These oscillations for very chiral swimmers also occur with a single population and are plotted in Fig. S2c. As the cluster rotates around the pole, the resulting vortex and torque on the drop also change sign (Fig. S2d) while remaining CW on average. This leads to the decrease in the mean late time value and the large standard deviation for high relative chirality in Fig. S2a. The exact value for the transition depends strongly on the details of the model, including in the near field. Importantly, however, this oscillating state in our system occurs for physically relevant values of the microscopic parameters.

#### B. Robustness to noise

We test the robustness of our system to noise both in orientation and translation, each assumed independent from the other, in Fig. S3.

A strong translational noise reduces the strength of collective effects, and in particular, the torque on the system $\tau$, as shown in Fig. S3. Increasing the rotational noise $D_r$ tends to delay the apparition of the vortex to higher values of the magnetic field, as shown in Fig. S3c. The peak value for the vortex strength is also smoothed. At high fields, increasing $D_r$ has no effect on the system as the torque from the magnetic fields dominates.

When considering values corresponding to the diffusion of a single magnetotactic bacteria in the bulk $D_r \sim$

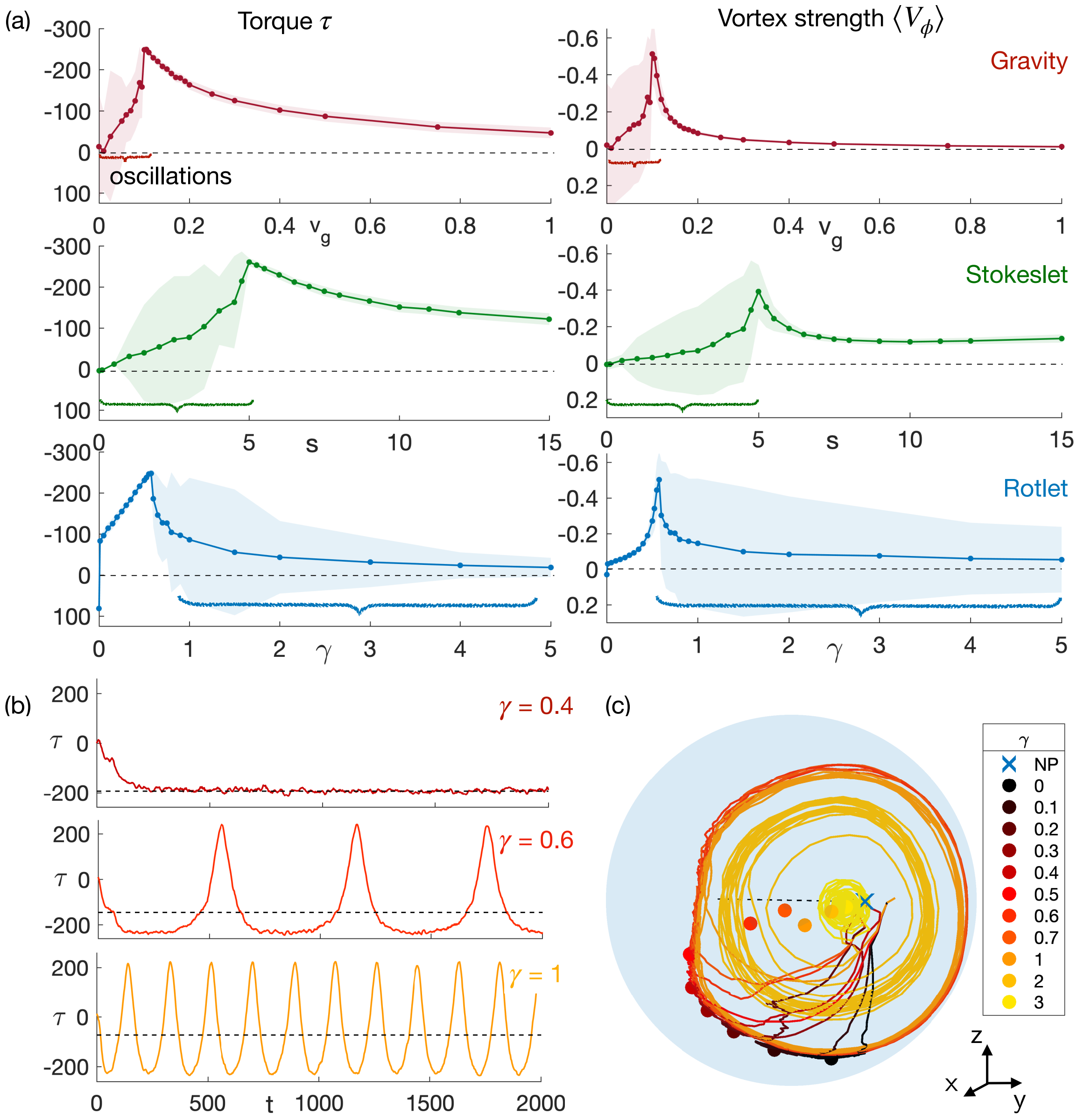


FIG. S2. Changing the microscopic parameters and transition to oscillations. (a) Torque on the drop (left) and vortex strength (right) when varying gravity $v_g$, Stokeslets $s$ and rotlets $\gamma$. In each case, there is an optimal value past which chirality becomes dominant and the long-term state is oscillatory, which leads to an increase in standard deviation outlined with an accolade. (b) Time variation of the torque on the drop due to 200 NS swimmers, for three values of chirality, $\gamma = 0.4, 0.6$ and 1, with the late time mean torque shown as a dashed line. For $\gamma = 0.6, 1$, the torque displays oscillations and changes direction. (c) Trajectory of the centre of a group of 200 NS swimmers with time for different values of $\gamma$, and onset of oscillations. The average late-time position of the cluster is shown as a dot.

$0.25 \mathrm{rad\,s}^{-1}$ [5], the influence of the rotational noise on the transition is small. However, in the suspension, we would expect contacts between swimmers to be an important source of noise in the system. While the corresponding values of $D_t$ and $D_r$ are hard to estimate, we show in Fig. S3 the effect of increased noise up to strengths much higher than the bulk ones. Overall, the qualitative properties of the system are very robust to noise, but we would expect the active noise in a suspension to affect quantitative values such as the strength of the field that maximises the rotation.

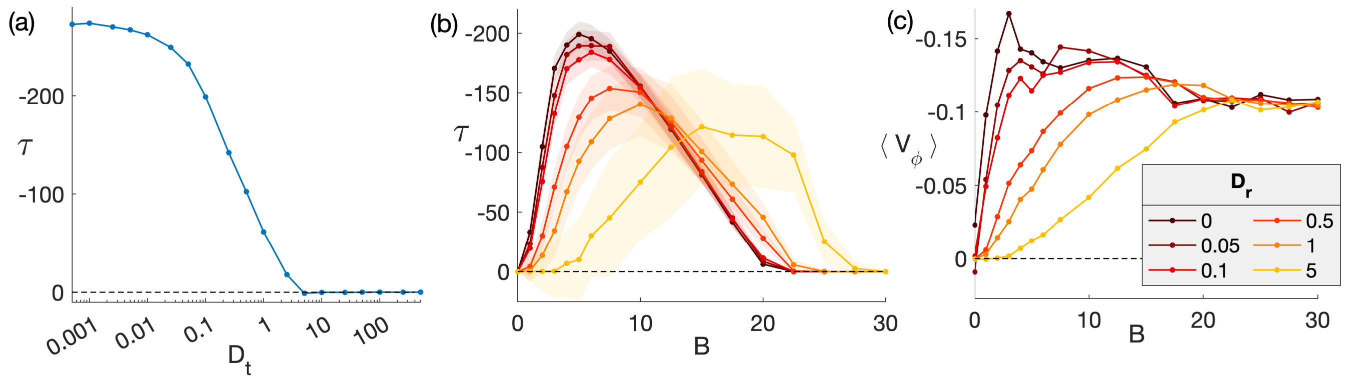


FIG. S3. (a) Variation of the strength of the torque $\tau$ on the drop with the translational noise strength $D_t$. (b) Variation of the torque $\tau$ on the drop with the magnetic field $B$ for different rotational noise strength $D_r$. The shaded areas show the standard deviation in time. The value of the rotational noise in the bulk is $D_r \approx 0.05$ in dimensionless units [5]. (c) The time-average of the azimuthal velocity $\langle V_\phi \rangle$ averaged over $r$ depending on the field $B$ for the same values of $D_r$.

#### III. IMAGES OF FORCE AND TORQUE DIPOLES INSIDE A NO-SLIP SPHERE

Here we derive expressions for the velocity and vorticity fields due to axisymmetric and transverse Stokeslets and rotlets inside of a rigid sphere. The derivation is based heavily on previous work carried out by previous authors, namely [8] for the axisymmetric Stokeslet, [9] for the transverse Stokeslet, [10] for the axisymmetric rotlet and [11] for the transverse rotlet.

Throughout this section we shall assume that the sphere is centred at the origin and has unit radius, that the viscosity $\mu = (8\pi)^{-1}$ and that the singularity is located at $\boldsymbol{x}_1 = \{0, 0, c\}$ in Cartesian coordinates with $0 < c < 1$ and has unit magnitude. The solution for more general parameters may then be obtained by applying rotations and scalings in conventional fashion.

Furthermore, we define the spherical polar coordinates

$$r = \sqrt{x^2 + y^2 + z^2}, \quad \theta = \cos^{-1}(z/r), \quad \phi = \tan^{-1}(y/x) \tag{12}$$

and the auxiliary quantities

$$\boldsymbol{x}_2 = \{0, 0, c^{-1}\}, \tag{13}$$
$$\boldsymbol{r}_1 = \boldsymbol{x} - \boldsymbol{x}_1, \tag{14}$$
$$\boldsymbol{r}_2 = \boldsymbol{x} - \boldsymbol{x}_2, \tag{15}$$
$$r_1 = |\boldsymbol{x} - \boldsymbol{x}_1| = \sqrt{r^2 + c^2 - 2rc\cos\theta}, \tag{16}$$
$$r_2 = |\boldsymbol{x} - \boldsymbol{x}_2| = \sqrt{r^2 + c^{-2} - 2rc^{-1}\cos\theta}. \tag{17}$$

In each case, the solution satisfies the boundary condition $\boldsymbol{u} = \boldsymbol{0}$ on the surface of the unit sphere and the Stokes equations within, except for the desired kind of singular behaviour at $\boldsymbol{x}_1$.

##### A. Axisymmetric Stokeslet

We consider a point force directed in the positive $z$-direction, $\boldsymbol{e}=\{0,0,1\}$. [8] give the solution in terms of singularities expressed as the Oseen tensor and its derivatives. Specifically,

$$\boldsymbol{u}\left(\boldsymbol{x}\right)=\boldsymbol{\mathcal{G}}\left(\boldsymbol{r}_1\right)\cdot\boldsymbol{e}+\frac{1-3c^2}{2c^3}\boldsymbol{\mathcal{G}}\left(\boldsymbol{r}_2\right)\cdot\boldsymbol{e}-\frac{1-c^2}{c^4}\left(\boldsymbol{e}\cdot\boldsymbol{\nabla}\right)\boldsymbol{\mathcal{G}}\left(\boldsymbol{r}_2\right)\cdot\boldsymbol{e}-\frac{\left(1-c^2\right)^2}{4c^5}\nabla^2\boldsymbol{\mathcal{G}}\left(\boldsymbol{r}_2\right)\cdot\boldsymbol{e}, \tag{18}$$

where the Oseen tensor is given by

$$\boldsymbol{\mathcal{G}}\left(\boldsymbol{r}\right)=\frac{\boldsymbol{I}}{|\boldsymbol{r}|}+\frac{\boldsymbol{rr}}{|\boldsymbol{r}|^3}. \tag{19}$$

Taking components and the curl respectively it is then straightforward (with appropriate software) to find components of the velocity and vorticity in spherical polar coordinates as follows.

$$\begin{aligned}u_r=\frac{1}{2}\Bigg(&2\cos\theta\left(\frac{(c-r\cos\theta)^2}{r_1^3}+\frac{1}{r_1}+\frac{\left(1-3c^2\right)\left(\left(\frac{1}{c}-r\cos\theta\right)^2+r_2^2\right)}{2c^3r_2^3}+\frac{\left(-1+c^2\right)^2\left(-3r^2+3r^2\cos(2\theta)+4r_2^2\right)}{4c^5r_2^5}\right.\\
&\left.+\frac{\left(-1+c^2\right)\left(-1+cr\cos\theta\right)\left(-3-3cr\cos\theta(-2+cr\cos\theta)+c^2r_2^2\right)}{c^7r_2^5}\right)+r\cos^2\phi\left(-\frac{2(c-r\cos\theta)}{r_1^3}\right.\\
&\left.+\frac{3-3c^2r_2^2+c^4\left(-3+5r_2^2\right)+cr\cos\theta\left(-9-6c\left(-1+c^2\right)r\cos\theta-3c^4\left(-1+r_2^2\right)+c^2\left(6+r_2^2\right)\right)}{c^6r_2^5}\right)\sin^2\theta\\
&+r\left(-\frac{2(c-r\cos\theta)}{r_1^3}\right.\\
&\left.+\frac{3-3c^2r_2^2+c^4\left(-3+5r_2^2\right)+cr\cos\theta\left(-9-6c\left(-1+c^2\right)r\cos\theta-3c^4\left(-1+r_2^2\right)+c^2\left(6+r_2^2\right)\right)}{c^6r_2^5}\right)\sin^2\phi\sin^2\theta\Bigg),\end{aligned} \tag{20}$$

$$\begin{aligned}u_\theta=&\frac{\left(-4c^9r_2^5-4c^7r_1^2r_2^5+r_1^3\left(12+c^2\left(-12+3\left(3-5c^2+2c^4\right)r^2+2r_2^2\left(-5+c^2\left(9-2c^2+\left(-1+3c^2\right)r_2^2\right)\right)\right)\right)\right)\sin\theta}{4c^7r_1^3r_2^5}\\
&+\frac{r\left(\left(2c^7r_2^5+r_1^3\left(-15+c^2\left(18+r_2^2-3c^2\left(1+r_2^2\right)\right)\right)\right)\sin(2\theta)-3c\left(-1+c^2\right)rr_1^3\sin(3\theta)\right)}{4c^6r_1^3r_2^5},\end{aligned} \tag{21}$$

$$u_\phi=0, \tag{22}$$

$$\omega_r=0, \tag{23}$$

$$\omega_\theta=0, \tag{24}$$

$$\begin{aligned}\omega_\phi=&\frac{r\left(\left(4c^7r_2^7+r_1^3\left(-15\left(-1+c^2\right)^2\left(1+c^2r^2\right)+3c^2\left(1-6c^2+5c^4\right)r_2^2+2c^4\left(1-3c^2\right)r_2^4\right)\right)\sin\theta\right)}{2c^7r_1^3r_2^7}\\
&-\frac{3\left(-1+c^2\right)r^2r_1^3\left(5+c^2\left(-5+2r_2^2\right)\right)\sin(2\theta)}{2c^6r_1^3r_2^7}.\end{aligned} \tag{25}$$

##### B. Transverse Stokeslet

For the transverse Stokeslet we assume that the unit force is oriented in the positive $x$-direction, $\boldsymbol{e} = \{1, 0, 0\}$. The solution by [9] is given in terms of hydrodynamic scalar potentials $\psi(r,\theta)$ and $\chi(r,\theta)$ from which the flow components are calculated as

$$u_r = -\frac{\cos\phi}{r^2}\frac{\partial}{\partial\theta}\left(\frac{1}{\sin\theta}\frac{\partial\psi}{\partial\theta}\right), \tag{26}$$

$$u_\theta = \frac{\cos\phi}{r}\frac{\partial}{\partial\theta}\left(\frac{1}{\sin\theta}\frac{\partial\psi}{\partial r}\right) + \frac{\cos\phi}{r\sin^2\theta}\chi, \tag{27}$$

$$u_\phi = -\frac{\sin\phi}{r\sin^2\theta}\frac{\partial\psi}{\partial r} - \frac{\sin\phi}{r}\frac{\partial}{\partial\theta}\left(\frac{\chi}{\sin\theta}\right). \tag{28}$$

The potentials themselves are given as follows (we note that our definition of $r_2$ differs from [9]).

$$\begin{aligned}\psi = {} & \frac{(r\cos\theta - c)r_1}{c} + \frac{(r-1)^2\left(\left(r\left(1-c^2\right)-2c^2\right)\cos\theta - 2c\right)}{2c} \\ & + \frac{4cr^2\cos^2\theta - \left(\left(c^2+1\right)r^2 + \left(5c^2+1\right)\right)r\cos\theta + c\left(\left(1-c^2\right)r^4 + \left(3c^2-1\right)r^2 + 2\right)}{2c^2 r_2},\end{aligned} \tag{29a}$$

$$\chi = \frac{2(r_1 - cr_2 - (r-1)(c\cos\theta + 1))}{c}. \tag{29b}$$

From this, we find the velocity components as

$$\begin{aligned}u_r = {} & \frac{1}{2}\cos\phi\Bigg(-\frac{2c(c - r\cos\theta)}{r_1^3} + \frac{4}{r_1} + \frac{1}{c^4 r_2^5}\left(3r\left(1+r^2+c^2\left(5+r^2\right)\right)\cos\theta\right. \\ & \left. + 3c\left(-2-\left(1+3c^2\right)r^2 + \left(-1+c^2\right)r^4 - 2r^2\cos(2\theta)\right) + 2c\left(1+r^2+c^2\left(5+r^2\right) - 8cr\cos\theta\right)r_2^2 - 8c^3 r_2^4\right)\Bigg)\sin\theta,\end{aligned} \tag{30a}$$

$$\begin{aligned}u_\theta = {} & \frac{\cos\phi\csc^2\theta}{4c^2 r r_2^3}\Bigg(-4 - 2\left(3+11c^2\right)r^2 + 4\left(-2+c^2\right)r^4 + \frac{r}{c}\left(\left(3+4r^2+c^2\left(19+6\left(2+c^2\right)r^2 - 2\left(-1+c^2\right)r^4\right)\right)\cos\theta\right. \\ & \left. - 2c\left(-1+c^2\right)r\left(-3+2r^2\right)\cos(2\theta) - \left(1+2r^2+c^2\left(5+4r^2\right)\right)\cos(3\theta) + 4cr\cos(4t)\right) \\ & + 2\left(1+5c^2+3\left(1+c^2\right)r^2 + 2cr\cos\theta\left(-5-3c^2+2\left(-1+c^2\right)r^2 + 2\cos(2\theta)\right)\right)r_2^2 - 8c^2 r_2^4 \\ & + \frac{3r}{c^2 r_2^2}\left(-2c\left(2+\left(3+8c^2\right)r^2 + 2r^4\right)\cos\theta\right. \\ & \left. + r\left(1+r^2+c^2\left(9+r^2\left(3+2r^2-2c^2\left(-3+r^2\right)\right)\right)\right) + \left(1+5c^2\right)\left(1+r^2\right)\cos(2t) - 2cr\cos(3\theta)\right)\sin^2\theta \\ & + \frac{2cr_2^3}{r_1^3}\left(r_1^2\left(-2\left(c^2+r^2\right) + 5cr\cos\theta - cr\cos(3\theta) + 3\left(-1+c^2\right)\left(-1+r^2\right)r_1 + 2r_1^2\right)\right. \\ & \left. + cr\left(-2\left(c^2+r^2\right)\cos\theta + cr(3+\cos(2t))\right)\sin^2\theta\right)\Bigg),\end{aligned} \tag{30b}$$

$$\begin{aligned}u_\phi = {} & -\frac{\csc^2\theta}{2c^2 r r_2^2}\Bigg(\frac{(cr - \cos\theta)}{cr_2}\left(r\left(1+r^2+c^2\left(5+r^2\right)\right)\cos\theta + c\left(-2-\left(1+3c^2\right)r^2 + \left(-1+c^2\right)r^4 - 2r^2\cos(2\theta)\right)\right) \\ & - \left(-6c^3 r + 4c\left(-1+c^2\right)r^3 + \left(1+5c^2+3\left(1+c^2\right)r^2\right)\cos\theta - 6cr\cos(2\theta)\right)r_2 \\ & - \frac{c}{r_1}\left(-2\left(c^2+r^2\right)\cos\theta + cr(1+3\cos(2\theta)) + \cos\theta r_1\left(3\left(-1+c^2\right)\left(-1+r^2\right) + 2r_1\right)\right)r_2^2 + 4c^2\cos\theta r_2^3\Bigg)\sin\phi.\end{aligned} \tag{30c}$$

The vorticity components are omitted due to their length.

##### C. Axisymmetric rotlet

For the axisymmetric rotlet we again consider an orientation in the positive $z$-direction, $\boldsymbol{e} = \{0, 0, 1\}$, corresponding to an anticlockwise rotation when viewed from $z = \infty$. [10] found that the solution can be written with the aid of just one additional image rotlet and reads

$$u_r = u_\theta = 0, \tag{31}$$

$$u_\phi = r\left(\frac{1}{r_1^3} - \frac{1}{c^3 r_2^3}\right)\sin\theta. \tag{32}$$

The vorticity is given by

$$\omega_r = \frac{2\cos\theta}{r_1^3} - \frac{3cr\sin^2\theta}{r_1^5} + \frac{-2c\cos\theta r_2^2 + 3r\sin^2\theta}{c^4 r_2^5}, \tag{33}$$

$$\omega_\theta = \left(\frac{3r(r - c\cos\theta)}{r_1^5} - \frac{2}{r_1^3} + \frac{3r(-cr + \cos\theta) + 2cr_2^2}{c^4 r_2^5}\right)\sin\theta, \tag{34}$$

$$\omega_\phi = 0. \tag{35}$$

##### D. Transverse rotlet

For the transverse rotlet, we assume the orientation is $\boldsymbol{e} = \{0,1,0\}$, so that the rotation is in the $x$-$z$-plane. The solution found by [11] is again in terms of potentials, much like the transverse Stokeslet found by [9], and the potentials are

$$\psi = -\frac{r_1}{c} + \frac{r}{c} - \cos\theta + \frac{r}{cr_2}\left(1 - r^2\right)\left(cr - \cos\theta\right) + \frac{1}{2}r\left(3 - r^2\right)\cos\theta + \frac{1}{2}\left(1 + r^2\right)\left(r_2 - \frac{1}{c}\right), \tag{36}$$

$$\chi = \frac{1}{r_1} - \frac{r\cos\theta}{cr_1} + \frac{\cos\theta}{c} - \frac{r}{c^2}\left(\frac{cr - \cos\theta}{r_2} + c\cos\theta\right). \tag{37}$$

Explicit expressions for the velocity and vorticity are again omitted here due to their extreme length.
